# Introducing the Y chromosome ancestral reference sequence - Improving the capture of human evolutionary information

**DOI:** 10.1101/2025.05.07.652589

**Authors:** Zehra Köksal, Annina Preussner, Jaakko Leinonen, Taru Tukiainen

## Abstract

Reference sequences are essential for reproducible genetic analyses but are often chosen without regard to evolutionary relevance within the analyzed species. The human Y chromosome (chrY) is widely used in evolutionary studies, yet current references represent evolutionarily young sequences, which can lead to misleading variant calling. To address this issue, we constructed a Y-chromosomal ancestral-like reference sequence (Y-ARS) to improve the detection of evolutionarily informative variants on the Y chromosome.

The Y-ARS was constructed by applying a weighted maximum parsimony approach to human and primate Y chromosome sequences. To benchmark the performance of the Y-ARS, 40 chrY short-read sequences from diverse haplogroups were aligned to Y-ARS and existing references (GRCh37, GRCh38, and T2T-CHM13). Overall, the Y-ARS yielded the highest and most consistent number of SNPs per sample (mean=1197; SD=105), while other references yielded on average fewer variants (mean=866–968) and showed greater variability across samples (SD=457–531) depending on their phylogenetic distance from the reference. Additionally, alignments to the Y-ARS resulted in calling solely SNPs with evolutionarily derived alleles, while alignments to other references resulted in calling on average 44% SNPs with ancestral alleles.

This study demonstrates how the existing reference sequences fail to capture the full range of evolutionary information on the chrY. The Y-ARS improves capturing evolutionary information on the chrY, making it a valuable resource for various evolutionary applications, such as TMRCA estimations and phylogenetic analyses. Finally, alongside the Y-ARS, we provide a publicly available tool, polaryzer, to annotate variants as ancestral or derived in pre-aligned chrY data.

**Significance statement:** Using current reference sequences results in calling genetic variants without information on whether the variant is ancestral or arose later in the species’ history, which is of interest for evolutionary studies. Here, we tackle this problem for the human Y chromosome by introducing the Y-chromosomal ancestral-like reference sequence, Y-ARS. The Y-ARS overcomes this issue by calling only evolutionarily derived variants on the Y chromosome in a reproducible way, which can directly be used for various downstream analyses in evolutionary genetics.

## Introduction

Reference sequences are essential for reproducible genetic analyses, as they establish a standard sequence and an universal coordinate system, enabling consistent gene annotation, functional analysis, and representation of genetic diversity in the form of genetic variants (Worley et al. 2017; Wong et al. 2020). The reference genomes of well-studied organisms, such as the human or mouse, are consistently improved by applying newer sequencing technologies and providing more complete references (Nurk et al. 2022). Whereas the classical reference sequences, such as human GRCh37 (GCF_000001405.25, NCBI Assembly), GRCh38 (GCF_000001405.40, NCBI Assembly) and T2T-CHM13 (T2T) (Nurk et al. 2022) are largely derived from only one contemporary sample, emerging pangenomes based on multiple samples contain more within-species diversity (Liao et al. 2023). However, all the current reference sequences lack information on evolutionary states of the variants, which is fundamental information for evolutionary studies.

In evolutionary and population genetic studies, the Y chromosome is one of the most widely utilized sequences. Since the MSY (male-specific region of Y chromosome) escapes recombination, all of its genetic variations are inherited together as a haplotype, making it a useful tool for tracing back mutational events for the study of demographic events. These Y-chromosomal haplotypes can be classified into haplogroups (named alphabetically from A to T) that share a common ancestor. Although haplogroup A is the oldest lineage, diverged from the rest at around 160-307 kya (Karmin et al. 2015; Hallast et al. 2023), Y chromosomes of the current human genome references are derived from evolutionarily young haplogroups R1b (GRCh37, GRCh38) and J1 (T2T), formed approximately 15-26 kya and 13-24 kya, respectively (Karmin et al. 2015; Hallast et al. 2023).

Using an evolutionarily young reference is problematic for evolutionary and population genetics studies. On the one hand, all alleles that are identical between the sample and the reference are disregarded as these emerge as non-polymorphic in variant calling, although some of these might contain evolutionarily relevant alleles. On the other hand, evolutionarily ancestral alleles might be called as sequence variants at sites where the reference contains a derived allele. As a result, the identified variants lack information on the allele polarization, i.e., whether the observed alleles are evolutionarily ancestral or derived. Instead, the called variants only represent differences between the sample and the reference genome. Yet, several applications, such as assessing sequence evolution, phylogenetic analyses, and demographic modeling, currently rely on knowledge of the evolutionary states of (Y-chromosomal) alleles (Kelleher et al. 2019; Speidel et al. 2019; Köksal et al. 2023).

For individual genomic sites, the ancestral states of nucleotides can be reconstructed using methods applying maximum parsimony (Collins et al. 1994) (e.g. MEGA (Tamura et al. 2021)) or maximum likelihood (Felsenstein 1981; Koshi & Goldstein 1996) (e.g. PAML and FastML (Ashkenazy et al. 2012)). For individual sites on the human Y chromosome, the ancestral states are usually reconstructed by comparing the samples to an evolutionary outgroup, such as primates (Karmin et al. 2015; Hallast et al. 2015). However, currently there is no easily applicable, streamlined and reproducible solution to aid in determining evolutionarily derived alleles on the human Y chromosome in a straightforward manner.

In this study, we constructed an ancestral-like reference sequence for the human Y chromosome (Y-ARS), to provide a reproducible solution to call only evolutionarily derived variants on the Y chromosome. To construct the Y-ARS, we applied weighted maximum parsimony (WMP) to determine ancestral alleles in the T2T Y chromosome sequence, using eight long-read human sequences of diverse major Y-chromosomal haplogroups for increased sequence overlaps and capture of human diversity. As evolutionary outgroups in the WMP, the four short-read primate sequences (chimpanzee, bonobo, gorilla, orangutan) were used. We benchmarked the Y-ARS with 40 short-read human sequences from all major haplogroups (A to T), which were aligned to the Y-ARS and existing references GRCh37, GRCh38 and T2T. To enable the use of Y-ARS beyond alignment-based applications (i.e., when handling VCF data), we developed polaryzer, a tool that annotates called variants from other references as ancestral or derived. Both, the novel Y-ARS reference sequence, as well as polaryzer, are made publicly available to simplify the process of determining evolutionarily derived alleles on the Y chromosome in a reproducible way.

## 1. Results

### The Y-ARS shows advantages in annotating ancestral states over SNP databases

To obtain the Y-chromosomal ancestral-like reference sequence (Y-ARS), we began assessing sites that are polymorphic among humans within the non-repetitive regions of the Y chromosome (∼15.6 million bp) (Fig. 1). At each polymorphic site (N=11,535 among the eight human sequences), we applied a weighted maximum parsimony approach (WMP) to infer the most probable ancestral allele using data from eight human and four primate Y chromosomes (Fig. 1). To gain the Y-ARS, we then converted all alleles identified as evolutionarily derived on the T2T Y chromosome (N=2,273) back to their ancestral state (Table S1). This number represents the SNPs that have accumulated on the T2T Y chromosome since its divergence from the most recent common ancestor (MRCA), i.e., the Y-ARS. This places the age of the Y-ARS at approximately 167 kya (CI 155-182 kya), which is in line with previous estimates dating the MRCA of the human Y chromosome to 160-307 kya (Karmin et al. 2015; Hallast et al. 2023).

**Fig. 1.**
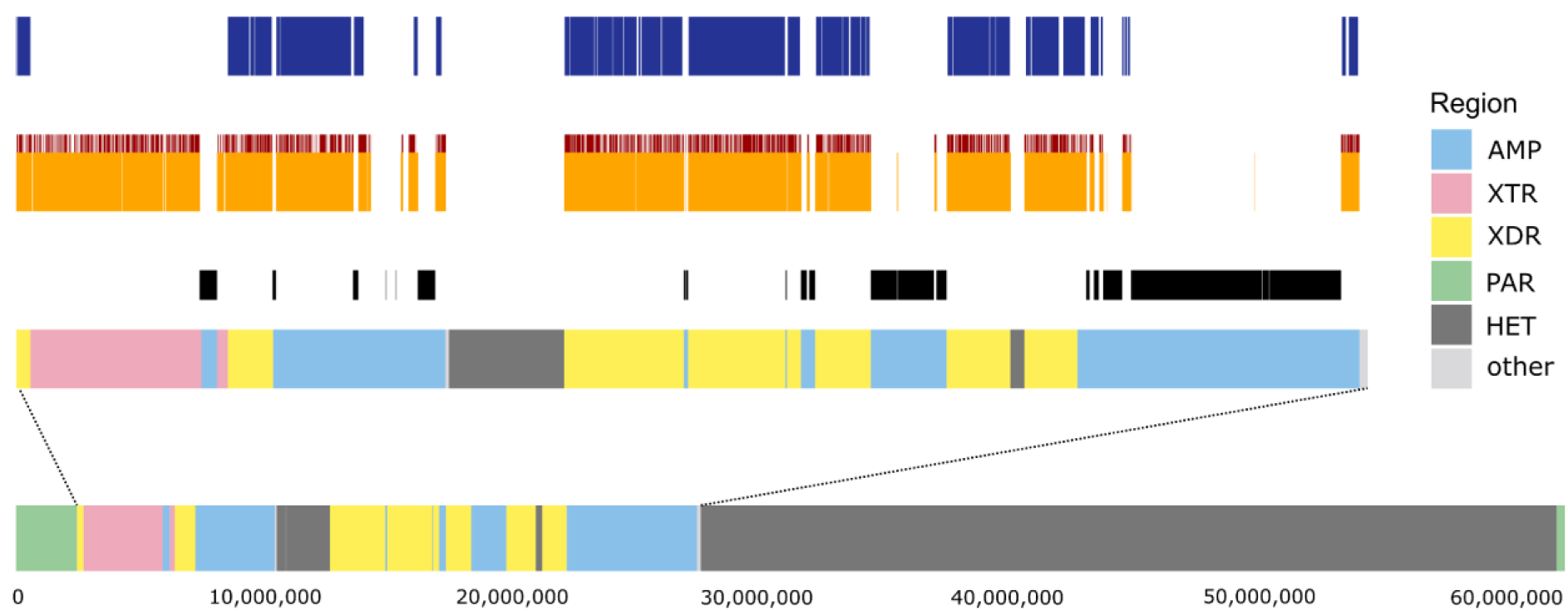
Y chromosome in T2T coordinates along with different Y chromosome sequence classes annotated: ampliconic (AMPL), X-transposed (XTR), X-degenerate (XDR), pseudoautosomal regions (PAR), heterochromatic (HET), other. The ancestral state reconstruction was done on 15.6 Mbp of XDR, XTR, and AMP regions excluding the highly repetitive regions (i.e., palindromes, TSPY and inverted repeats) marked in black. The locations of the N=11,535 polymorphic sites utilized in the ancestral state reconstruction are marked with orange along with the N=2,273 derived sites identified on the T2T sequence marked with red. Poznik et al. (2013) regions accessible for short-read data mapping (10.5 Mbp) are marked with dark blue.

To validate the ancestral states of the alleles in the Y-ARS, we accessed SNP entries in the human Y-SNP database YBrowse (https://ybrowse.org/). YBrowse is a comprehensive database of haplogroup-defining Y-chromosomal SNPs, which includes information on the ancestral and derived alleles of each SNP submitted by individual researchers. The ancestral alleles of SNPs in the YBrowse database were compared with the allele in the Y-ARS of the corresponding Y-chromosomal sites. Along the Y-chromosomal area of interest (15.6 Mb), we identified in total 1,455,104 sites with an existing YBrowse annotation. These corresponded to 1,563,189 SNP entries, given some sites had several database entries. For the vast majority of the identified sites (99.6%; N=1,448,650), the allele observed in the Y-ARS corresponded to the ancestral allele in the YBrowse database as expected, supporting the validity of the ancestral state of the Y-ARS alleles.

However, for a small fraction of the sites (0.6%; N=9,032), the allele in the Y-ARS carried the derived allele (i.e., a contradiction) according to the YBrowse database. Yet, the vast majority of these sites (N=7,492) were not polymorphic among our data, possibly characterizing (sub)haplogroups not represented by the eight samples used to reconstruct the Y-ARS in the current study. These could be possible errors in the database caused by allele switch-ups between the ancestral and derived allele. For the sites that were polymorphic among our data (N=1,540), we assessed these contradictions in more detail (Fig. S1; Fig. S2). A considerable fraction of these contradicting annotations (N=1,218) occurred at sites with multiple annotations with different alleles reported as the ancestral allele (Fig. S2A). Here, back-mutations (i.e., two mutational events at the same genomic site occurring by chance) appeared to account for many of the overlapping annotations (Fig. S2Aa). Some of the sites with multiple database entries were not necessarily explainable by back-mutations, but showed support for the observed Y-ARS allele being ancestral (Fig. S2Ab, S2Ac). A small portion of the sites with contradicting annotations did not have multiple annotations, but were annotated only as derived in the database (N=322). Yet, on all of these sites our data showed support for the Y-ARS allele more likely to be ancestral than derived (Fig. S2B). For a small number of sites (N=33) we could not infer the ancestral alleles with high confidence, due to high mutability (Fig. S2Bg) or high rates of missingness (Fig. S2Bd,e). Altogether, examining these contradicting annotations showed that the Y-ARS provided more often reliable annotations for Y chromosome SNPs than the YBrowse database on the compared sites, which could be driven by the fact that the YBrowse database also contains submissions mainly from individual studies or testing groups, rather than comprehensive population-wide data or phylogenetic reconstructions.

Although the majority of alleles on the Y-ARS appeared to be confidently ancestral, a key challenge in ancestral state reconstruction remains in accurately determining ancestral alleles in samples that are phylogenetically close to the focal node. (i.e., haplogroup A0). As the focal node is located right before the A0 haplogroup, it becomes difficult to infer whether the allele carried by A0 has formed before or after the A0 split from the most recent common ancestor of the human Y chromosome. To evaluate this possible haplogroup A bias within the Y-ARS, we further extracted all SNPs, where an allele was carried exclusively by the sample of haplogroup A0 among humans. We identified 2,520 of such sites (Table S2), which we further explored in additional human sequences of haplogroup A0 and primate data (see Material and Methods). While the majority of alleles at these sites (86%; N=2,165) supported the Y-ARS allele being the ancestral allele, all sites could not be conclusively re-evaluated, resulting in the true ancestral allele remaining ambiguous on 355 sites, listed in Table S3.

### Benchmarking the Y-ARS using short-read data underlines its advantages in calling a balanced number of evolutionarily derived SNPs

To quantify how the Y-ARS impacts variant calling on the Y chromosome, we aligned 40 short-read human Y chromosome sequences from all major haplogroups (A-T) to the Y-ARS and existing references GRCh37, GRCh38 and T2T. After quality filtering and excluding poorly mappable regions (Fig. 1; Fig. S3), the Y-ARS resulted in calling the highest number of SNPs when averaged across all samples (N=1,197 normalized SNP count per 10 Mbp), compared to GRCh37 (N=866), GRCh38 (N=878) and T2T (N=968) (Fig. 2A). Furthermore, the number of SNPs called upon alignment to the Y-ARS were relatively constant across samples (standard deviation SD=105; range=1029-1585), while the GRCh37 (SD=505; range=206-2774), GRCh38 (SD=531; range=129-2821), T2T (SD=457; range=383-2765) displayed more variation in the total number of SNPs called (Levene’s test *p*=0.016) (Fig. 2A,B).

**Fig. 2.**
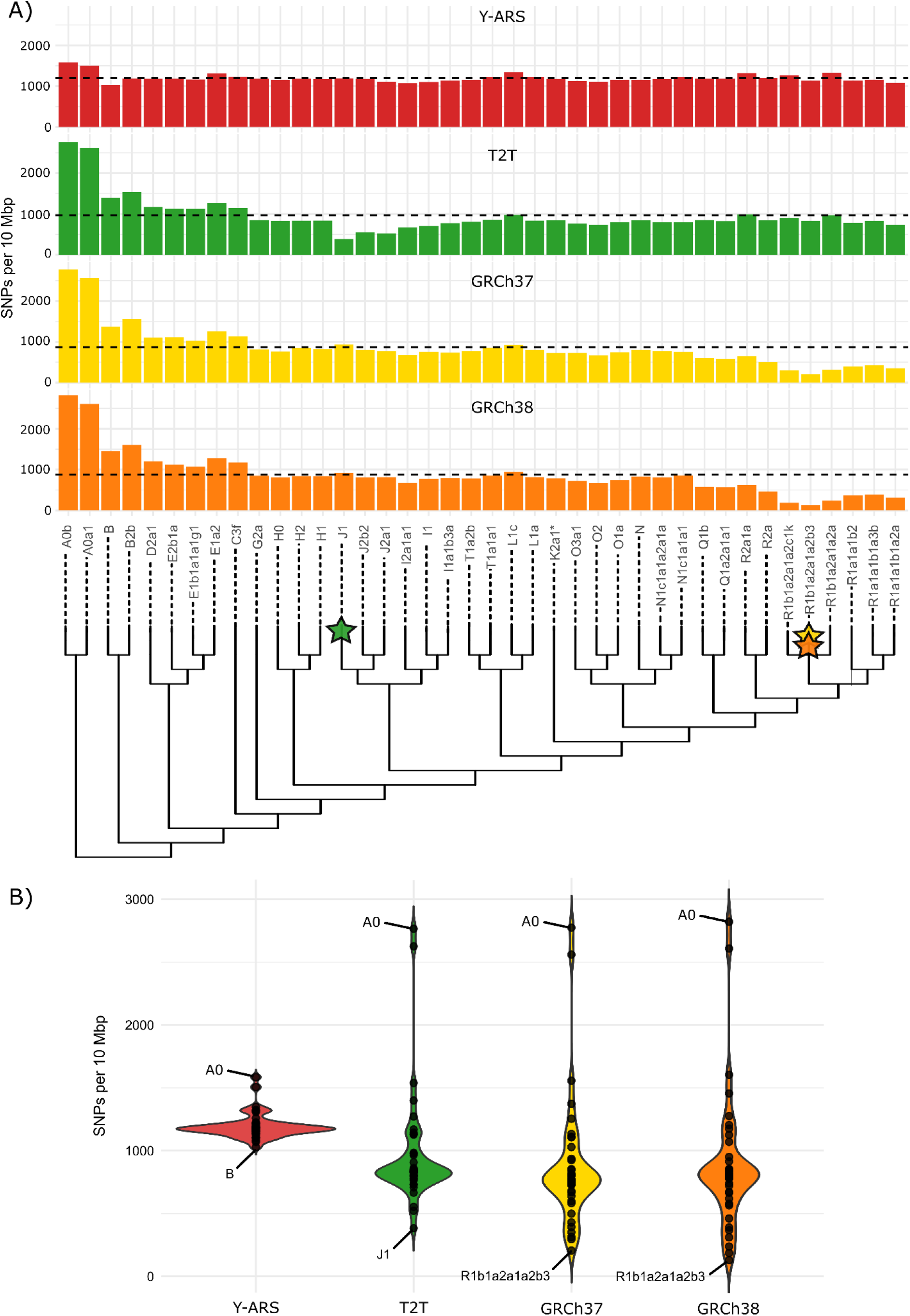
**A)** Number of chrY SNPs normalized by per-sample coverage (and multiplied by 10 million) per reference across all samples. The different colors correspond to different reference sequences used for alignment. Phylogenetic location of references T2T, GRCh37 and GRCh38 representing downstream haplogroups are marked with stars on the phylogenetic tree. References GRCh37 and GRCh38 represent haplogroup R1b1a1b1a1 and T2T represents haplogroup J1a2a1a2c1a1. B) Variability in SNP counts on each reference sequence for all samples. Samples with the lowest and highest number of SNPs observed for each reference are annotated.

For the three standard reference sequences, the number of SNPs called was lower in cases where the reference sequence and the sample sequence belonged to phylogenetically close haplogroups (Table S4; Fig. 2A,B). The T2T reference, which represents a haplogroup J1 sequence, resulted in calling the smallest number of variants for sample HG01253 of haplogroup J1 (N=383) compared to other samples aligned to the same reference (N=968 mean; N=836 median across all samples). Similarly, alignments to references GRCh37 and GRCh38, which both represent haplogroup R1b sequences, resulted in calling fewer variants for samples of haplogroup R1b compared to samples of other haplogroups. For instance, sample NA19652 of haplogroup R1b1a2a1a2b3 resulted in calling six times fewer SNPs on GRCh38 (N=129) compared to other samples aligned to the same reference (N=878 mean; N=808 median across all samples). Accordingly, the number of SNPs increased when the sample and the reference belonged to phylogenetically distant haplogroups. Generally, most SNPs were called for the samples among the oldest and most basal branches. Samples of haplogroup A0 carried up to 3 times more SNPs when aligned to GRCh37, GRCh38 and T2T, and 1.3 times more SNPs when aligned to the Y-ARS compared to other samples aligned to the same references (Fig. 2).

### Evolutionarily young references result in calling a mixture of ancestral and derived alleles

Since the Y-ARS represents an ancestral human Y chromosome, all SNPs called upon alignment to Y-ARS are expected to be evolutionarily derived. However, since references GRCh37, GRCh38, and T2T are evolutionarily young haplogroups, samples aligned to these result in calling SNPs, which can be from an evolutionary perspective derived or ancestral. To determine the evolutionary state (i.e. the polarization) of alleles upon alignment to GRCh37, GRCh38 and T2T, we annotated alleles that match with the Y-ARS allele as ancestral and alleles that deviate from the Y-ARS as derived, using a custom-made tool “polaryzer” (accessible on Github). When polarizing the variants among the 40 samples, we first lifted alignments from GRCh37 and GRCh38 to T2T coordinates to minimize the effect of losing variants during liftover (Fig. S4).

When considering the evolutionary states of the called variants upon alignment to GRCh37, GRCh38, and T2T, on average 44% of the called SNPs represented evolutionarily ancestral alleles (Fig. 3A). The average proportions were fairly similar across these references: 40% on GRCh37 (N=216) (range=9-49%; 18-830 SNPs), 46% on GRCh38 (N=206) (range=11-54%; 11-776 SNPs) and 47% on T2T (N=360) (range=36-52%; 86-1218 SNPs) (Fig. 3A, Fig. S5). The remaining 56% of the called SNPs represented evolutionarily derived alleles on these references. In contrast, all SNPs called on Y-ARS-aligned samples represented evolutionarily derived variants, formed after the MRCA of the human Y chromosome (Fig. 3).

**Fig. 3.**
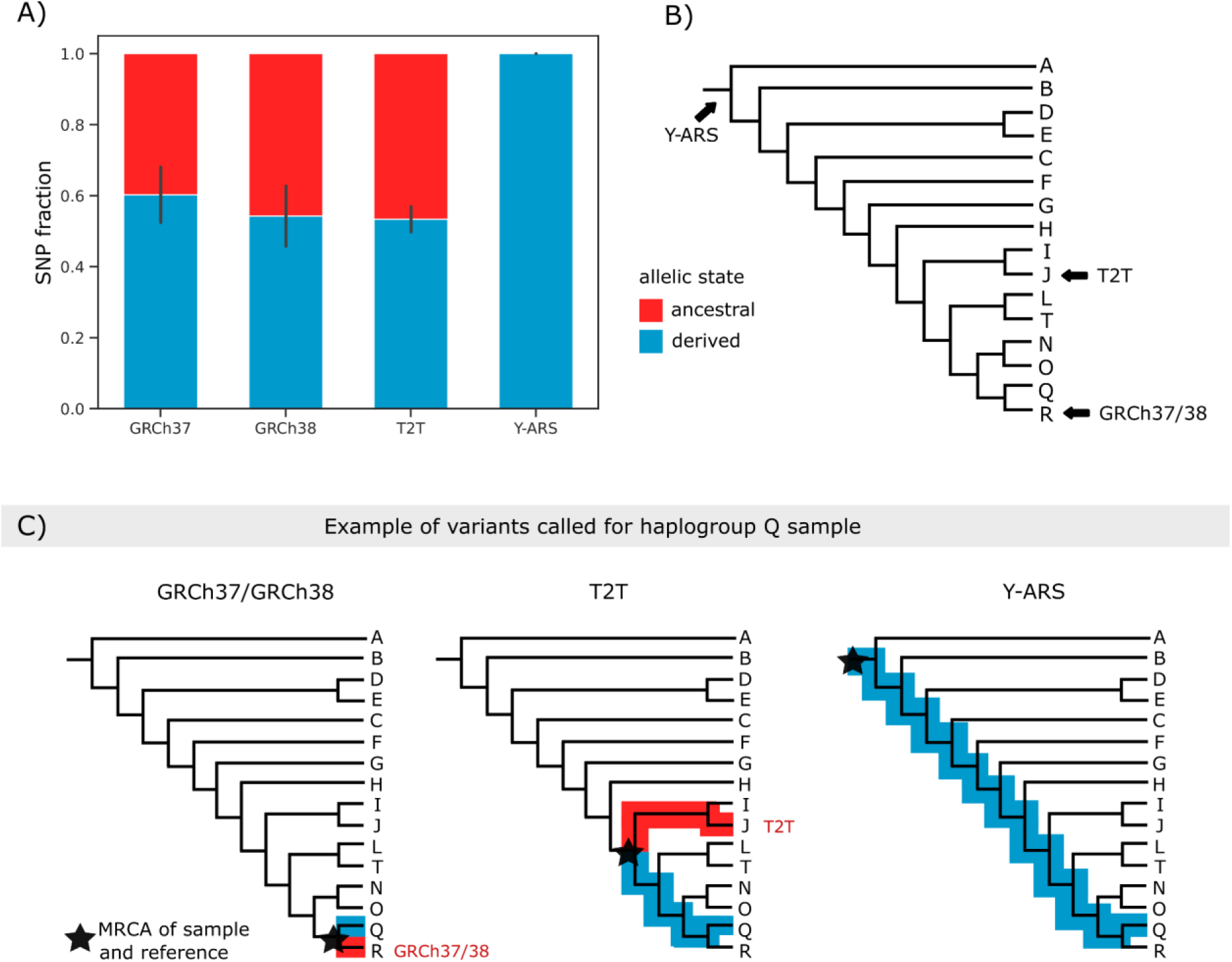
**A)** Fraction of evolutionarily derived (blue) and ancestral (red) alleles of variants observed on average over all samples from software polaryzer. The barplot height represents averages over all samples and the error bars the standard deviations. B) Phylogenetic location of each reference sequence. C) Example breakdown of SNPs captured for haplogroup Q sample upon alignment to references GRCh37/GRCh38, T2T, and Y-ARS, divided into evolutionarily derived in the reference (red) and evolutionary derived in the sample (blue). Variants that are evolutionarily derived in the reference appear as ancestral alleles in variant calling, as illustrated in panel A. Only SNPs formed after the most recent common ancestor (MRCA) of the sample and the reference are captured.

To better characterize these observed SNPs on each of the references beyond the ancestral and derived states, we assessed how many of these had an existing annotation in SNP databases (dbSNP, ISOGG, YFull). The dbSNP-annotated sites represent known genetic variants across populations, while the ISOGG and YFull annotations highlight phylogenetically informative SNPs on the Y chromosome, which are commonly used in haplogroup classification and Y chromosome phylogenetic analyses. Thus, sites that lack these annotations likely lack a phylogenetic context, or are private or rare SNPs in the sample or the reference.

When aligning to GRCh37, GRCh38 and T2T, on average 84% of the called SNPs had an existing annotation (48% in dbSNP, 69% in ISOGG/YFull) (Fig. S6). Upon alignment to the Y-ARS, the percentage of annotated variants was 94% (88% in dbSNP; 72% in ISOGG/YFull) (Fig. S6). This was expected given the greater phylogenetic distance between the samples and the Y-ARS compared to other references, which facilitated the detection of SNPs that arose earlier in the phylogenetic tree. The proportion of haplogroup-defining alleles on Y-ARS-aligned samples (average 72%) was similar to that of the other references (average 69%). However, samples aligned to the Y-ARS carried 1.9-fold more haplogroup-defining alleles, since all evolutionarily derived alleles were present in the sample (rather than in the reference sequence) (Fig. S7).

When examining whether the proportion of annotated variants differed across samples aligned to the same reference, all haplogroup R1b samples (HG01785, NA19652, and HG00096) were outliers for references GRCh37 and GRCh38 (Fig. S7). These samples showed a notably lower proportion of annotated SNPs upon alignment to GRCh37 and GRCh38 (average 33%) compared to other samples aligned to the same references (average 83%) (Fig. S6; Fig. S7). This trend was expected, as a more recent split between the haplogroups of the sample and reference allows for fewer mutations to accumulate in the sample that are absent in the reference. In these cases, private mutations (i.e., non-annotated variants) accounted for a large fraction of the total variants. Intriguingly, haplogroup J1 sample (HG01253) did not show a similar decrease in the proportion of annotated SNPs when aligned to the T2T reference (average 72%) compared to other samples aligned to the same reference (average 86%) (Fig. S6; Fig. S7). This could be explained by the larger phylogenetic distance between sample J1 and the T2T reference compared to R1b samples and the GRCh37 and GRCh38 references. This was supported by the total number of SNPs observed, as HG01253 (haplogroup J1) and T2T differed in N=383 SNPs, while NA19652 (haplogroup R1b1a2a1a2b3) and GRCh38 differed in N=129 SNPs (Fig. 2A).

As expected, the alignment to the Y-ARS resulted in calling only a few haplogroup-defining SNPs per sample with ancestral alleles (median N=5). Interestingly, two samples of haplogroup A0 showed an enrichment for ancestral alleles of haplogroup-defining SNPs (N=122 and N=134) (Fig. S7). In these haplogroup A0 samples, these SNPs were mostly characteristic for haplogroup A1, as annotated in the ISOGG and YFull databases. As we described before, these haplogroup A1-defining SNPs likely contain annotation errors for haplogroup A1 rather than wrongly reconstructed alleles on the Y-ARS (see example in Fig. S2Bh) deriving from the close proximity of haplogroup A to the focal node. The challenges in determining the true haplogroup-defining allele (i.e., the derived allele) may result in a higher likelihood of annotation errors as seen in the database (Fig. S1).

## 2. Discussion

Although current reference sequences, such as GRCh37 and GRCh38 have both been used for more than a decade as a standard tool in genetic research, the effect of their evolutionary state on variant calling has not been considered thoroughly. While some efforts have been made to improve the study of mitochondrial DNA (Behar et al. 2012), the Y chromosome remains largely unassessed in this context. To overcome this issue, we constructed an ancestral-like reference sequence for the Y chromosome (Y-ARS), using weighted maximum parsimony for SNPs on ∼15.6 Mbp of non-repetitive regions of the Y chromosome. To benchmark the use of the Y-ARS as a reference sequence, we further aligned 40 short-read sequencing samples to the Y-ARS and existing references GRCh37, GRCh38 and T2T.

Our validations of the Y-ARS showed that the number of sites (N=2,273) at which we reverted the T2T template allele to the ancestral state, were in accordance with expectations given an average Y-chromosomal mutation rate. When evaluating the Y-ARS alleles using the comprehensive database of Y-chromosomal SNPs, YBrowse, we observed that the majority (99.6%) of sites on the Y-ARS were supported by existing database information, as expected. Only for a small portion of sites on the Y-ARS (0.4%), we observed contradicting database annotations, which were in most cases providing a more reliable annotation in our data compared to the database. These contradictions could be explained for instance by back-mutations or mislabeling of reference and ancestral allele. The latter was particularly noticeable for SNPs defining haplogroup R, which is the haplogroup of the GRCh37 and GRCh38. Although the database information was intended to validate the ancestral states of alleles in the Y-ARS, the Y-ARS turned out to provide more reliable ancestral annotations on several sites. The strength of the ancestral state reconstruction approach is that it relies on comprehensive phylogenetic assessment and systematic cross-species comparison, while YBrowse annotations originate mostly from independent submissions from specific study cohorts or genetic testing groups, with limited standardization or evaluation of the submissions. These findings strongly underline that caution is needed when relying solely on any database annotations. The Y-ARS provides additional advantages in determining the allele polarization of SNPs not yet reported in databases with a phylogenetic context.

Although the Y-ARS showed high confidence for the majority of alleles to be ancestral, a few sites could not be annotated confidently. These included sites with only a few data points, highly mutable sites, and sites with haplogroup A0-specific alleles, for which the location close to the focal node complicated the ancestral state reconstruction. Even when including additional data from samples of haplogroup A0, the ancestral state remained ambiguous for sites that did not show variation beyond this haplogroup. Thus, it is possible that the Y-ARS could contain alleles that are in fact specific to haplogroup A (Table S2). In particular, this may be reflected in the increased total number of variants called for haplogroup A0 samples upon alignment to the Y-ARS (N>1500), compared to the average number of SNPs observed across all samples when aligning to the Y-ARS (N=1197). Nevertheless, this slight increase in SNPs could also be attributed to the fact that haplogroup A0 is the most basal Y-chromosome lineage, having diverged from all other haplogroups approximately 160-307 kya (Karmin et al. 2015; Hallast et al. 2023), thereby undergoing a longer period of branch-specific evolution.

When evaluating the Y-ARS in Y chromosome sequence alignment and variant calling, it displayed advantages compared to existing references. Aligning samples to the Y-ARS resulted in consistent SNP calling across different haplogroups (range=1029-1585) of contemporary sequences, due to calling all SNPs formed since the MRCA of the Y chromosome. Samples aligned to the other references (GRCh37, GRCh38, T2T), showed more variability in the number of SNPs called (range=129-2821), depending on the phylogenetic location of the sample in comparison to the reference. This variation results from calling SNPs formed after the split of the reference and sample haplogroups, while evolutionarily derived SNPs emerged before the split appear as monomorphic. The most drastic decrease in the number of SNPs (up to 6-fold) was observed for haplogroup R1b samples when aligning these to GRCh37 and GRCh38, since these samples and references share a very recent common ancestor (7-17 kya estimated from our data).

In addition to providing consistent SNP calling across haplogroups, the Y-ARS resulted in calling only evolutionarily derived SNPs, while the other references resulted in calling a mixture of evolutionarily derived (54% on average) and ancestral (46% on average) variants. These evolutionarily ancestral variants reflect sites where the reference sequence carries a derived allele, and as such are redundant for analyses focused on the sample of interest. A similar pattern was also evident in the number of known haplogroup-defining variants identified in the sample, which was nearly twice as high when using the Y-ARS reference (70%) compared to other references (37%). We observed that the number of haplogroup-informative variants decreased (down to 3-21%) in particular for samples carrying haplogroup R1b upon alignment to GRCh37 and GRCh38, due to the increased capture of private or rare mutations for these samples. This highlights that aligning samples to a phylogenetically close reference sequence results not only in calling fewer variants, but also significantly decreases the proportion of phylogenetically informative variants captured. Given that haplogroup R1b is relatively common across Europe (Myres et al. 2011), the loss of information could be concerning for variant studies on Y chromosome data of European origin when aligning to GRCh37 or GRCh38.

While we emphasize the strengths of the Y-ARS for Y chromosome sequence alignment and ancestral state annotation, it is important to acknowledge its current limitations. At present, the ancestral states annotated on the Y-ARS are limited to single nucleotide variations among the 15.6 Mb of non-repetitive DNA sequence. These are the most commonly used genetic variants in Y chromosome research, yet they represent only a subset of the potential genomic variations. For studies focusing on (the evolution of) structural variants, it could be worthwhile in the future to examine an ancestral sequence that also includes indels and structural variants (outside of palindromic, repetitive regions), or a pangenome, although the repetitive nature of the Y chromosome might impede these attempts.

In essence, the Y-ARS enables calling of solely evolutionarily derived variants, whereas alternative references demand manual variant annotation, which is time-consuming, error-prone and lacks a standardized pipeline. Understanding the evolutionary state of Y-chromosomal variants is crucial for several downstream analyses and applications in Y chromosome research, including for instance sequence age estimation (Solé-Morata et al. 2017; Poznik et al. 2013), refined phylogenetic classification of variants and haplogroups (Köksal et al. 2024; Hallast et al. 2015) and demographic modeling (Sahakyan et al. 2021; Batini et al. 2015). Particularly, when investigating novel SNPs to increase the resolution of the Y chromosome haplogroups, identifying the evolutionarily derived SNPs, their phylogenetic hierarchy and geographic distribution can shed light on potentially unexplored prehistoric and historic migration pathways.

Overall, this new reference sequence represents the estimated ancestral states for Y-chromosomal SNPs on the most recent common ancestor of humans, allowing for consistent variant calling across different haplogroups. The Y-ARS can be seamlessly integrated into existing pipelines of Y chromosome sequence alignment. To enhance the broader usability of the Y-ARS, we also offer a tool, polaryzer, to annotate the evolutionary state of alleles on pre-existing VCF files. In conclusion, the new Y-ARS provides a consistent and reproducible way for identifying evolutionary derived variants on the human Y chromosome.

## 3. Conclusion

This study demonstrates how current references (GRCh37, GRCh38, T2T) fail to capture the full range of evolutionary information on the chrY. The Y-ARS improves capturing evolutionarily derived sites, making it a valuable resource for various evolutionary genetics applications, such as TMRCA estimation, phylogenetic analyses, and demographic modeling. We further provide a publicly available tool, polaryzer that utilizes the information in the Y-ARS to determine whether SNPs reported in VCF files (after alignment to GRCh37, GRCh38, T2T) are ancestral or derived. Further, we share the resource of generating this Y-ARS, to be extended on other species as well.

## 4. Materials and Methods

### Ancestral state reconstruction

In the following, we aimed to approximate the Y-ARS - the most recent common ancestor of the human Y chromosome sequences - by reconstructing the ancestral state at each Y-chromosomal site (Fig. 4). To reconstruct the ancestral states at a node of interest (i.e., focal node) it is necessary to combine (A) alleles at the same genomic position of the investigated samples, (B) the samples’ phylogenetic relationships, and (C) apply an evolutionary model suitable for the data.

**Fig. 4.**
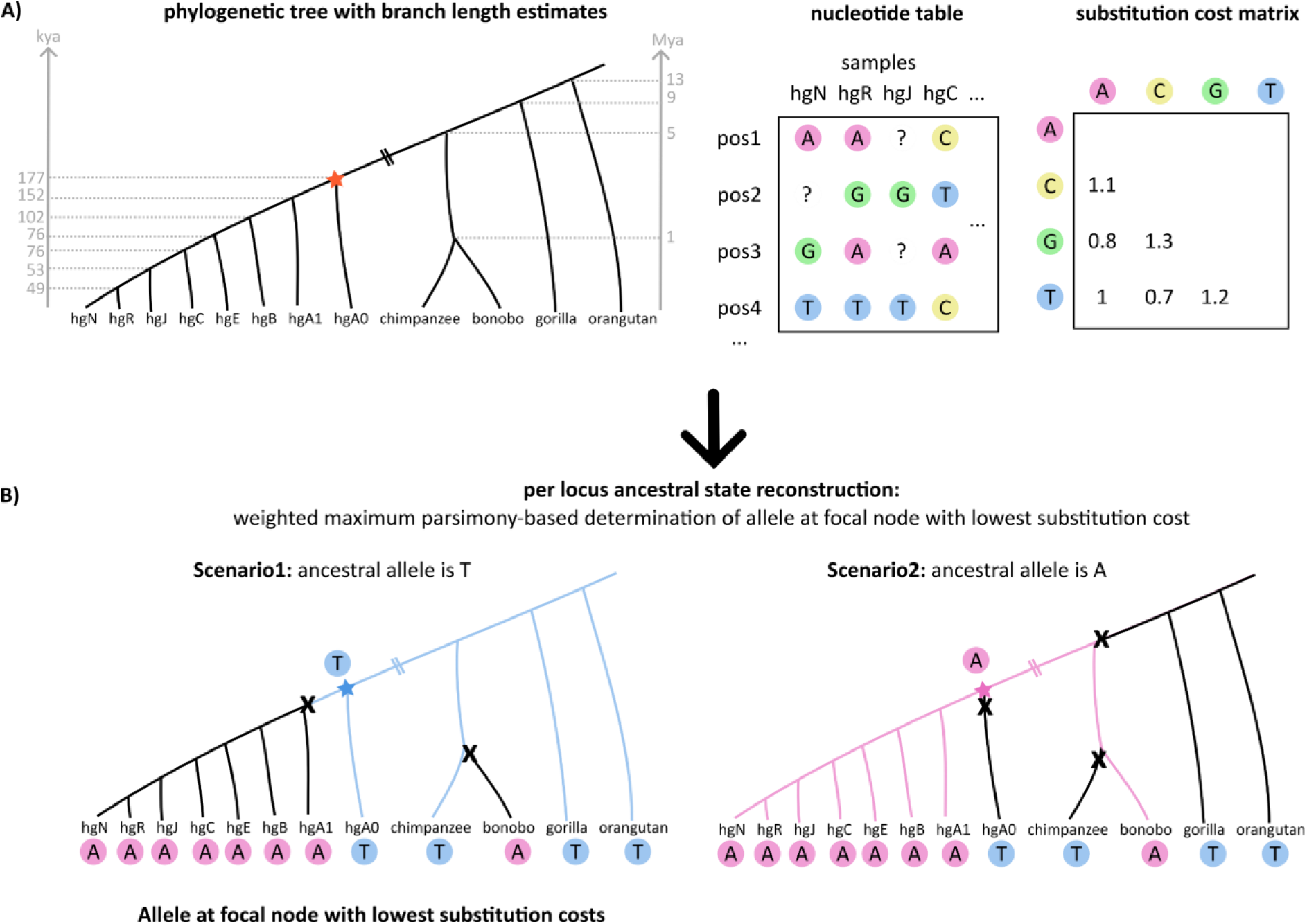
Weighted maximum parsimony approach for ancestral state reconstruction per locus. (A, left) Phylogenetic relationships of orangutan, gorilla, bonobo, chimpanzee and the human haplogroups A0, A1, B, C, E, J (T2T-CHM13), N, R. The focal node, i.e. most common ancestor of humans, is marked with a star. Lineage split times are presented in gray in thousand (kya) and million years (Mya). Other inputs include the (A, middle) nucleotide table of all polymorphic sites and (A, right) defined nucleotide substitution costs. (B) An example for allele cost calculation with alleles T (scenario 1) and A (scenario 2) as potential focal node alleles, where the costs are calculated by dividing the nucleotide substitution cost by the branch length factor. The scenario with the lowest total cost (here allele T) is the estimated ancestral allele at the focal node.

### Identifying the variable sites among Y chromosome sequences

To generate the Y-chromosomal ancestral reference sequence (Y-ARS), we selected the T2T-CHM13-v.2.0 Y chromosome sequence (T2T-Y) (haplogroup J1a2a1a2c1a1) as a template to maintain an already established coordinate system. Ancestral states were reconstructed at the sites along the T2T-Y template sequence, using long-read HiFi sequencing data of seven human samples of haplogroups A0b, A1a, B2b, C1a, E1b, N1a, R1b (Table S5). The samples were processed according to the long-read sequencing data processing pipeline shown on Github (see https://github.com/ZehraKoksal/Y-ARS/tree/main/Commands). In short, all FASTQ files per sample were merged, and then aligned to the T2T reference using minimap2 (-ax map-hifi -p 0.95 --secondary=yes -N 1 -a -L -eqx --MD) (Li 2018). Resulting BAM files were sorted using samtools v1.18 (Li et al. 2009), and variants were called only for the Y chromosome using freebayes v0.9.21 (-p 1 --min-mapping-quality 50 --min-base-quality 20 -- no-indels --no-mnps --no-complex) (Garrison & Marth 2012). After identifying polymorphic positions within all samples, these 14,528 target positions were re-called for all samples (-- report-monomorphic). The variants were filtered by keeping only sites with phred-scaled quality score QUAL>=20 within X-degenerate regions (XDR), X-transposed regions (XTR) and ampliconic regions (AMP). We removed loci located within inverted repeats (IRs), arms of the palindrome regions 1-8 (P1-8), and the testis specific protein Y-linked (TSPY) repeat arrays (Rhie et al. 2022) (Table S6,S7), which yielded a total of 11,535 polymorphic sites used for ancestral state reconstruction.

In the next step, we assessed 9,395 out of the 11,535 sites polymorphic among the human sequences in primate sequences (2,140 loci were excluded due to their location on the human-specific XTR). Two short-read great ape sequences from chimpanzee, bonobo, gorilla and orangutan (Table S5) were downloaded from Sudmant et al. 2013. After trimming of the FASTQ files using trimmomatic v0.39 (Bolger et al. 2014), the reads were aligned to the T2T reference sequence using bwa v07.7.17 mem (Li & Durbin 2009), sorted and merged using samtools. Upon addition of read groups, PCR duplicates were removed using picard MarkDuplicate v2.26.4 (Broad Institute 2019). The resulting BAM files were indexed using samtools and base quality scores were recalibrated using Genome Analysis Toolkit (GATK) v4.2.0.0 BaseRecalibrator and ApplyBQSR (Auwera & O’Connor 2020). Variable and non-variable alleles were called at the 9,395 pre-defined sites using BCFtools v1.9 (Li 2011) commands mpileup, call and filter to call SNPs with minimum mapping quality ≥ 50, minimum base quality ≥ 20 and read depth > 2 (bcftools mpileup -Ou -q 50 -Q 20 | bcftools call --ploidy 1). Details on the commands used for data processing are shared on Github.

The two samples of each primate species were combined to one representative sample in preparation for the ancestral state reconstruction. For this, (1) alleles shared by both individuals were kept; (2) if one individual had a missing allele, the other individual’s allele was kept; (3) if the individuals had different alleles, this locus was characterized as missing (N). Next, we combined the allelic states for the four primate species with the allelic states of seven human samples (and the T2T-Y reference), encompassing altogether 11,535 Y-chromosomal sites used for ancestral state reconstruction.

### Phylogenetic relationships of samples and substitution cost matrix

The phylogenetic tree of the samples was generated using the known relationships of these samples (Fig. S8). We added lineage divergence time estimates for the human samples from Hallast et al. (2023) and for the primates from (Kivell 2019). To avoid a bias in favouring mutational events on longer branches, the branch lengths were normalized to the shortest branch having a scaled length of 1. This was done by taking the decadic logarithm of the branch age estimates and increasing this value by 6. The phylogenetic tree is presented in Fig. S8. We further incorporated a simple nucleotide substitution cost matrix in the ancestral state reconstruction, which is a modified substitution model of the Kimura-2-parameter model (Kimura 1980) with symmetrical and ranked substitution costs favouring transitions over transversions.

### Weighted maximum parsimony approach for reconstructing ancestral states

To construct the Y-ARS, we used a weighted maximum parsimony approach to estimate ancestral alleles among the 11,535 loci. We combined the information of the alleles observed along the samples’ sequences (nucleotide matrix) together with the phylogenetic tree including branch lengths and a substitution cost matrix (Fig. 3). Using this information, we created a custom python script (shared on Github) to calculate the most parsimonious allele for the focal node, i.e., the most recent common ancestor of humans. At each locus, the allele with the lowest mutational costs at the focal node was selected as the most parsimonious allele (i.e., most likely ancestral allele).

### The ancestral-like Y chromosome reference sequence (Y-ARS)

After determining the most likely ancestral alleles, we converted all derived SNPs on the T2T-Y FASTA sequence back to ancestral states (N=2,273) using the SeqIO module from the Bio package v1.84 in Python v3.11.11. We replaced the Y chromosome in the T2T reference with the Y-ARS, keeping all other chromosomes in the reference unaltered.

To estimate the age of the Y-ARS (i.e. focal node), we used a rate of 8.71× 10^−10^ (CI 9.43×10^−10^ - 8.03× 10^−10^) mutations per position per year (Helgason et al. 2015), number of positions at non-repetitive regions of the Y chromosome (15,588,924 bp), and the number of derived sites on the T2T sequence (N=2,273).

### Annotating the Y-ARS using YBrowse information

To assess if the converted SNPs match with existing database annotations, we downloaded haplogroup annotations from the Human Y Chromosome Pangenome Browser “YBrowse” (https://ybrowse.org/; version update 14.02.2025), which compiles information from major Y chromosome databases as well as single submissions by researchers. The downloaded SNPs were filtered by removing SNPs with unknown YCC or ISOGG haplogroup annotations and SNPs annotated at position 1 or at chromosomes other than “chrY”, yielding in 1,783,937 SNPs with a haplogroup annotation (including sites with multiple annotations).

### Examining the Y-ARS for haplogroup A0 bias

Due to the unique location of the A0 branch relative to the most recent common ancestor of all humans (i.e. the focal node) in the phylogenetic tree, SNPs exclusive to haplogroup A0 might be biased in the absence of any primate data despite a thorough ancestral state reconstruction. To investigate for bias towards possible SNPs exclusive to haplogroup A0, we further investigated the 2,520 positions with an allele specific to A0 among all human samples by accessing these positions in two of the 40 human short-read sequencing data of haplogroup A0 (HG01890 and HG02982) (Table S8) and all six short-read primate sequences (Table S5). The alleles at the positions of interest were extracted from the generated BAM files using BCFtools (bcftools mpileup -Ou -q 50 -Q 20 -R 2520_sites.csv -f T2T.fasta sample.bam | bcftools call --ploidy 1 -O v -o out_sites.vcf -c ; bcftools filter -e ‘TYPE=“INDEL“’ out_sites.vcf -o sites_noindels.vcf). Additionally, the Y chromosome FASTA sequences of the three primate reference sequences, PanTro3 (NC_072422.2, NHGRI_mPanTro3-v2.0), PanTro6 (NC_006492.4, Clint_PTRv2) and PanPan1 (NC_073273.2, NHGRI_mPanPan1-v2.0_pri) were accessed and aligned to the T2T reference sequence using minimap2. In the sorted and indexed BAM file, bcftools mpileup was used to extract the alleles at defined sites using the above command. The commands are shared in the accompanying Github repository.

### Benchmarking the Y-ARS using 40 short-read sequences across different haplogroups

To evaluate the performance of the Y-ARS, we assessed the alignment and variant calling of 40 short-read sequencing male samples from the 1000 Genomes Project (1000 Genomes Project Consortium et al. 2015; Fairley et al. 2020) with this new reference. The samples were representative of all major haplogroup clades from A0 to T (Table S7). We utilized paired-end data of low coverage WGS from the GRCh38 data collection. The short-read sequences were processed according to the pipeline on Github, as described in short for primate data in section “Identifying the variable sites among Y chromosome sequences”. After processing the FASTQ files, all sequences were aligned to the Y-ARS, and existing references GRCh37, GRCh38, T2T-CHM13-v.2.0 with BCFtools. The variants were filtered to exclude indels and sites with depth<2 and sites falling outside the well-mappable area on the Y chromosome (Poznik et al. 2013). To compare the called positions, all coordinates were lifted to T2T with R-package rtracklayer (Lawrence et al. 2009) in R v.4.1.2 (R Core Team 2021). We used ISOGG (v.15.73) (https://isogg.org/tree/) and YFull (v10.01) (https://www.yfull.com/tree/) haplogroup annotations for annotating the sample-specific SNPs.

### Normalization of SNP counts and age estimation

Since the samples had differing coverages (Fig. S8), we normalized the SNP counts to compare these across samples by dividing the raw SNP count by coverage and multiplying this by 10,000,000 to estimate the SNP counts per 10 million base pairs.

We further calculated the age of split for each sample from the reference sequences: GRCh37, GRCh38 and T2T by the following equation;

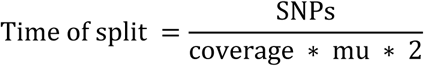

where the SNPs correspond to the raw number of SNPs and mu to the mutation rate 8.71× 10^−10^ (CI 9.43× 10^−10^ - 8.03× 10^−10^) mutations per position per year (Helgason et al. 2015), and coverage to the number of positions with at least 2x coverage on Y chromosome region of interest (10.5 Mb). For the Y-ARS we did not split the number of SNPs by two, since the variants captured represent SNPs captured in one sequence (i.e., the sample).

### Determining ancestral and derived alleles in VCF files using polaryzer

Polaryzer v1.0 (https://github.com/ZehraKoksal/Y-ARS/tree/main/Polaryzer) was run using the VCF files of all 40 samples (generated using variant calling thresholds: read depth ≥2 and excluding indels). The VCF files generated upon alignment to GRCh37 and GRCh38 were lifted over to reference T2T using rtracklayer in R v.4.1.2 and then used as input for polaryzer after filtered for non-repetitive regions. The specific commands used are available in the Github repository. To illustrate the fraction of ancestral and derived alleles per sample and reference sequence, barplots were generated using seaborn v0.13.2 in python v3.11.11.

## Data availability

All 1000 Genomes Project population samples used in the analyses are available from the NHGRI Repository at Coriell. Other sequences are available at the European Nucleotide Archive (long-read human: PRJEB58376, primate: PRJNA189439). Code for generating the Y-ARS is available on Github (https://github.com/ZehraKoksal/Y-ARS). The polaryzer tool for determining the allelic state of variants reported in VCF files is available on Github (https://github.com/ZehraKoksal/Y-ARS/tree/main/Polaryzer). The Y-ARS sequence will be available in Zenodo after acceptance to the journal.

## Acknowledgments

This work was supported by a Postdoctoral Fellowship from Linköping University (to Z.K.), research funding from the Doctoral Programme of Population Health at the University of Helsinki (to A.P.), Research Council of Finland (grant numbers 315589 and 320129 to T.T.) and the HiLIFE Fellows Program (T.T.).

We thank Leonor Gusmão and Vania Pereira for discussions and advice generating an artificial ancestral reference sequence, Mikko Rautiainen for discussing the details of alignment of sequencing data, Wolfram Höps for his advice on genome assemblies (and NAHRs), and Pille Hallast for sharing her expertise on human and primate Y chromosome data.

