## Supplementary Figures for "Introducing the Y chromosome ancestral reference sequence - Improving the capture of human evolutionary information"

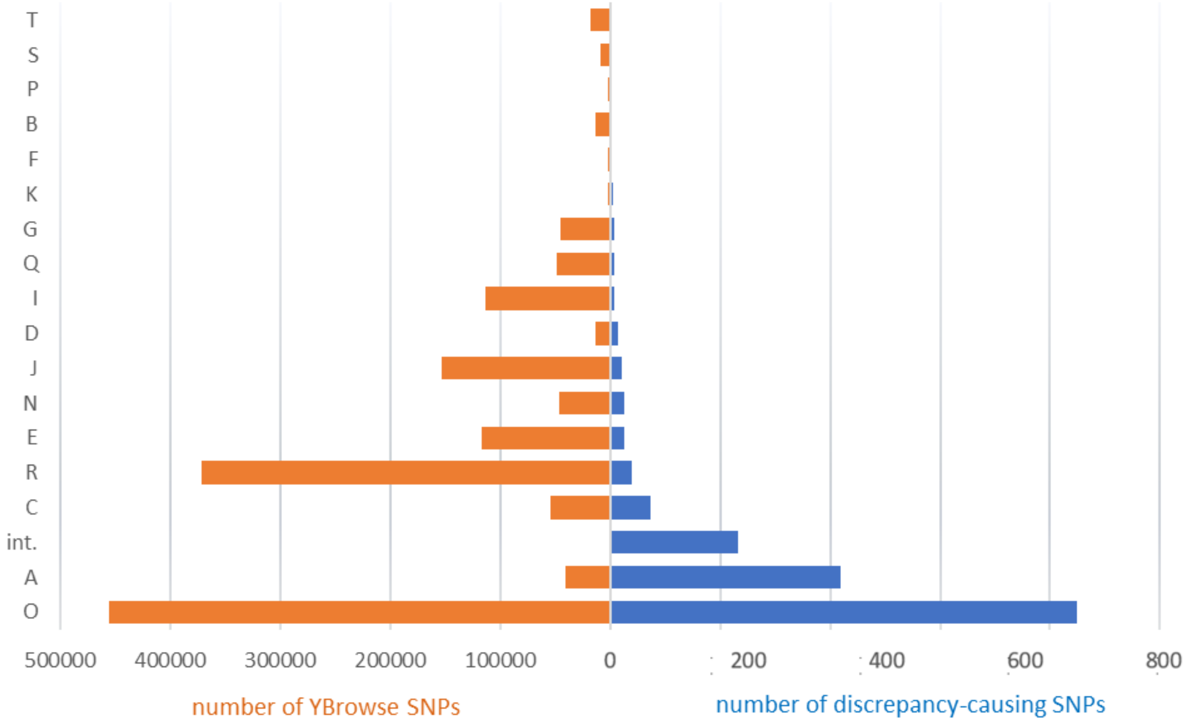


**Fig. S1** Number of haplogroup-defining SNPs per haplogroup in YBrowse (orange) (total=1,499,841) and number of SNPs causing discrepancies with ancestral alleles defined in Y-ARS sequence (blue) (total=1,396 SNPs). Annotation "int." comprise intermediate haplogroups BT, CT, CF. Most discrepancies were found for haplogroup O defining SNPs, which has the most SNP entries in the database. Second-most discrepancies are caused by haplogroup A, plausibly indicating inaccurate entries in the database due to the location of the haplogroup close to the focal node.

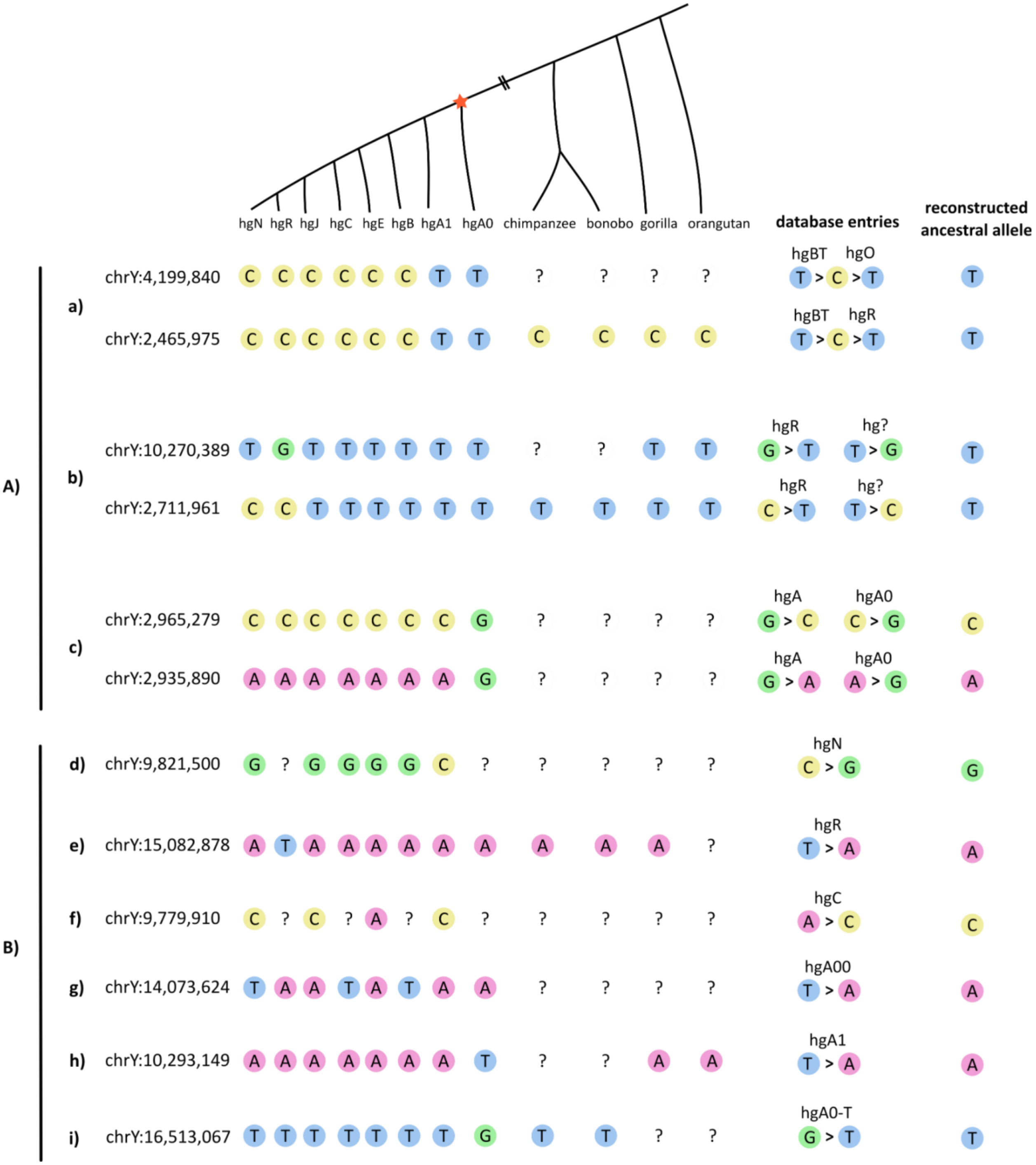


**Fig. S2** Examples of observed alleles among human and primate samples at Y-chromosomal loci with contradicting YBrowse database annotations compared to the reconstructed ancestral allele. A) Cases where the database contains multiple entries, that can be explained by the data. B)

Cases where the YBrowse derived allele does not match the Y-ARS allele. The contradictions between the database and the reconstructed ancestral alleles can be due to (a) back-mutations, (b, e) wrong database entries that likely resulted from using reference sequences of downstream haplogroups, (c, d, f) discrepancies in upstream haplogroups or inaccurate database entries, and (g) high mutability at genetic loci. The remaining examples (h, i) contain loci where the haplogroup A0 carries an allele distinct from other samples.

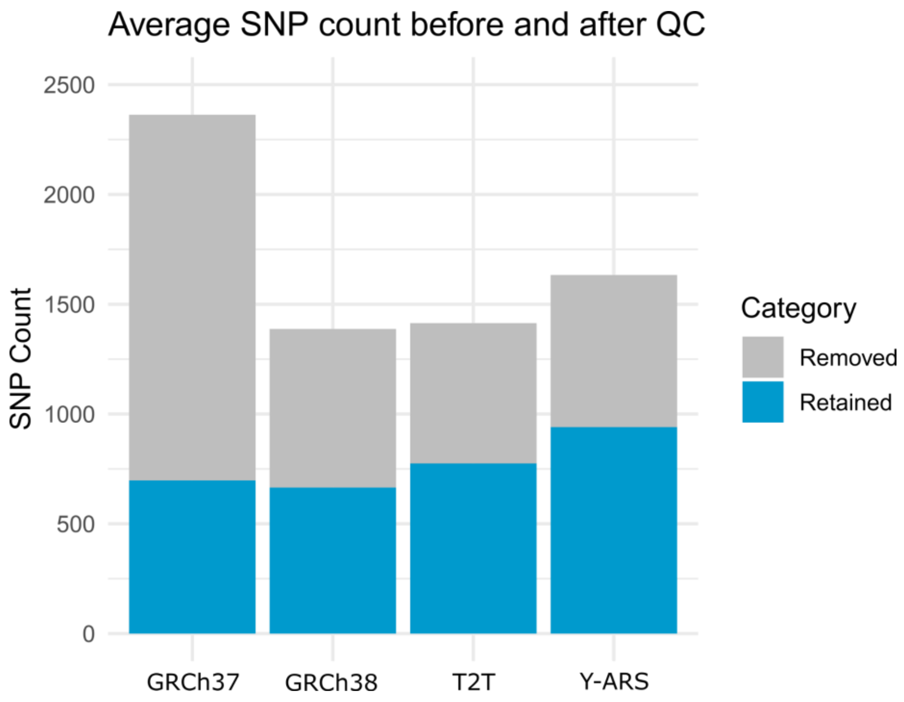


**Fig. S3** Average SNP count per reference before and after quality control. Blue indicates retained SNPs after quality control (QC) that are within non-repetitive regions defined by Poznik et al. (2013) accessible for short read data mapping, while gray indicates removed sites mapping outside these regions. Before quality control, GRCh37 alignments resulted in calling the highest number of SNPs on average across samples caused by excess read mapping on non-informative regions of the Y chromosome, such the centromere and Yq heterochromatin arm.

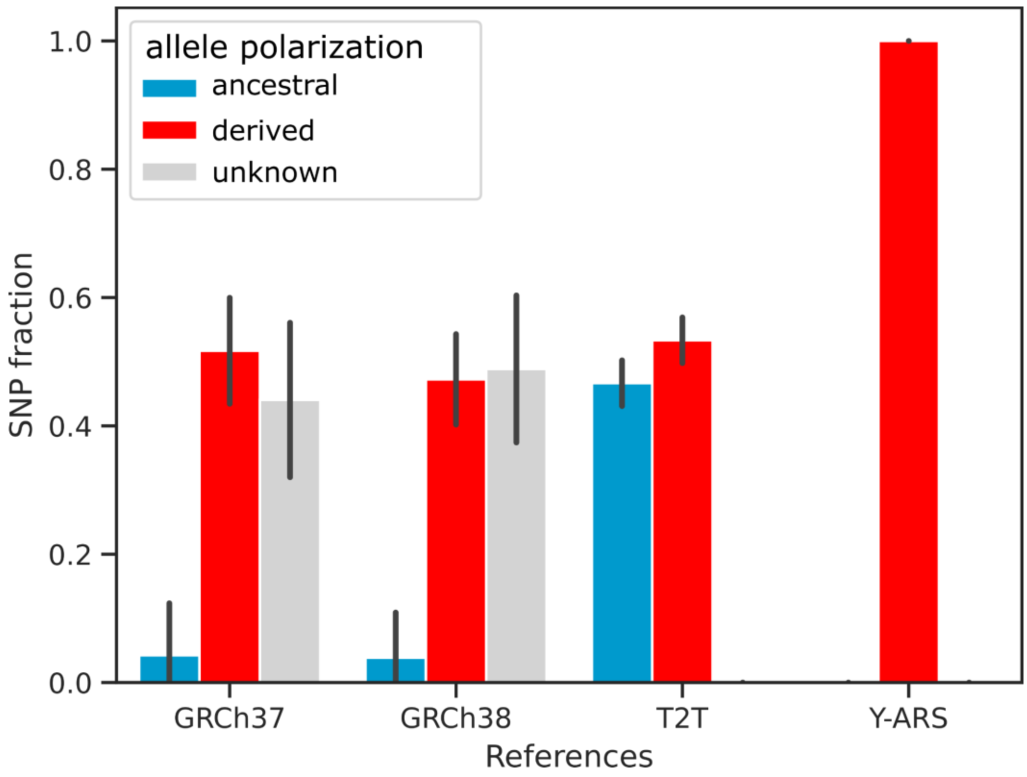


**Fig. S4** Fraction of SNPs annotated as ancestral (blue), derived (red) and unknown (grey) on each reference using the software polaryzer when running polaryzer to VCF files that are not lifted to T2T coordinates. The barplot height represents averages over all 40 samples and the error bars the standard deviation. Polarizing variants on VCF files in GRCh37/GRCh38 coordinates results in a high proportion of alleles without an annotation (44%-49%), caused by lifting over variants from Y-ARS (T2T coordinates) to GRCh37 or GRCh38 to generate dictionary files for polaryzer. Thus, better results can be obtained when lifting over GRCh37/GRCh38 coordinates to T2T prior to using polaryzer, as a liftover the this direction (GRCh37/GRCh38->T2T) seems more successful than in the opposite direction for sites carrying the ancestral alleles.

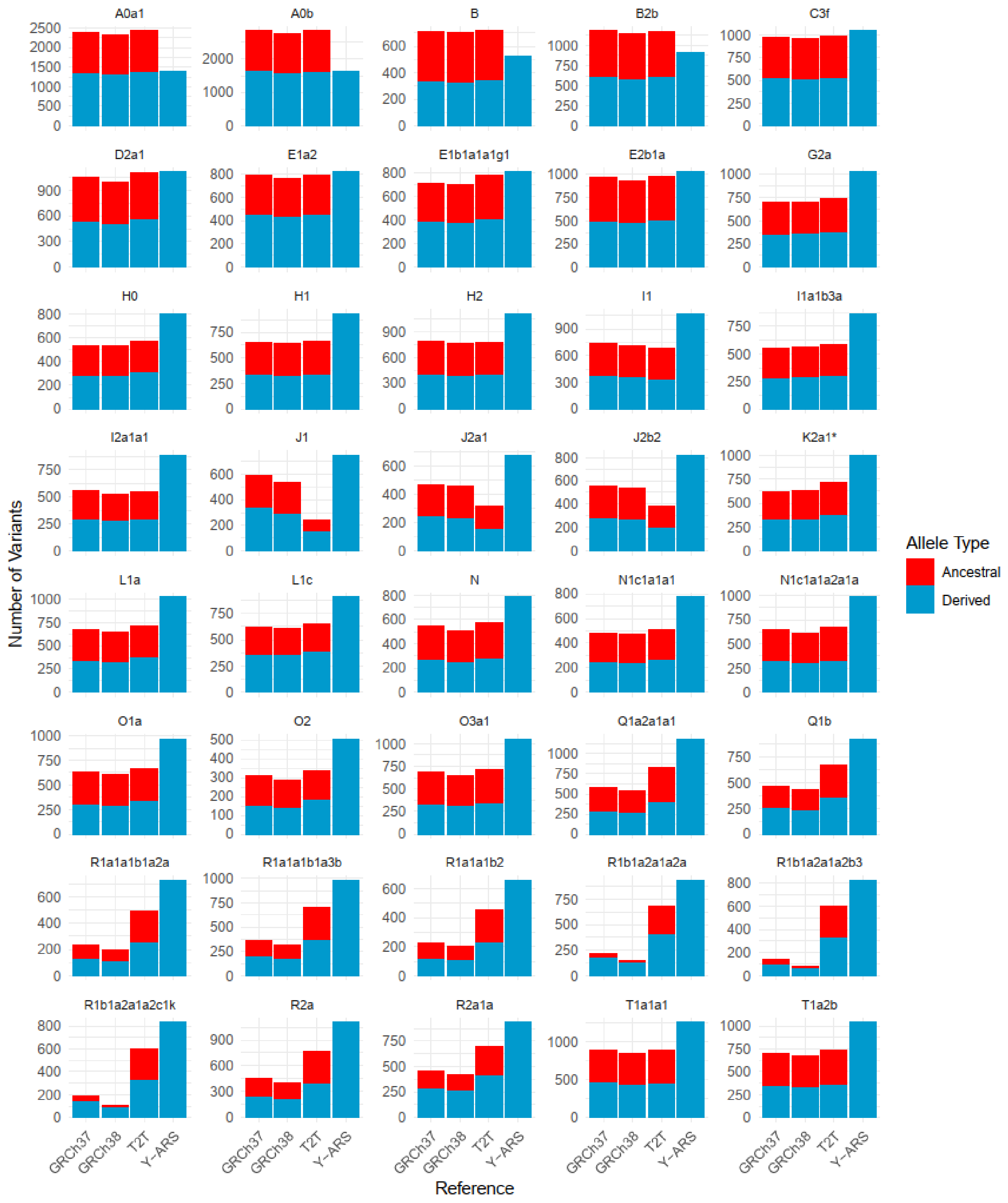


**Fig. S5** Number of variants of ancestral and derived allelic states across all 40 samples and all 4 references.

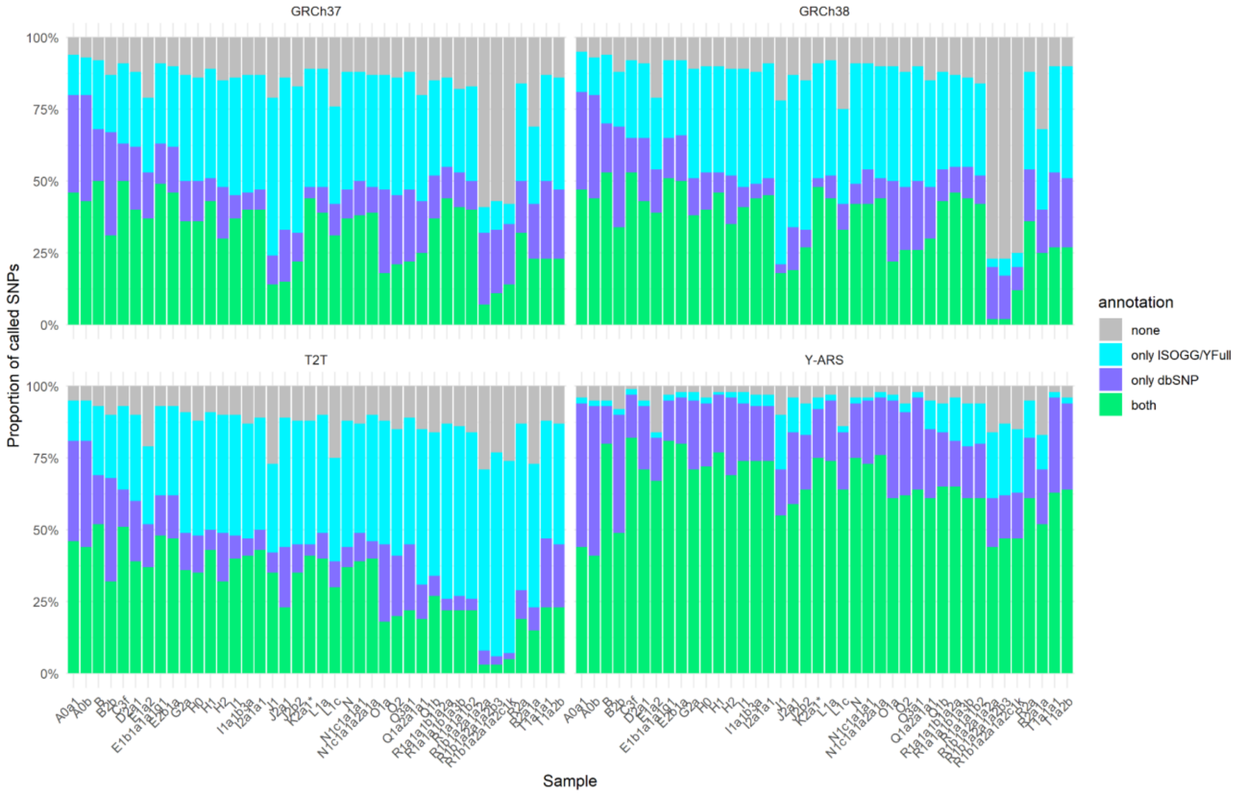


**Fig. S6** Proportion of SNPs have an annotation in dbSNP or ISOGG/YFull databases across the 40 samples and each of the references.

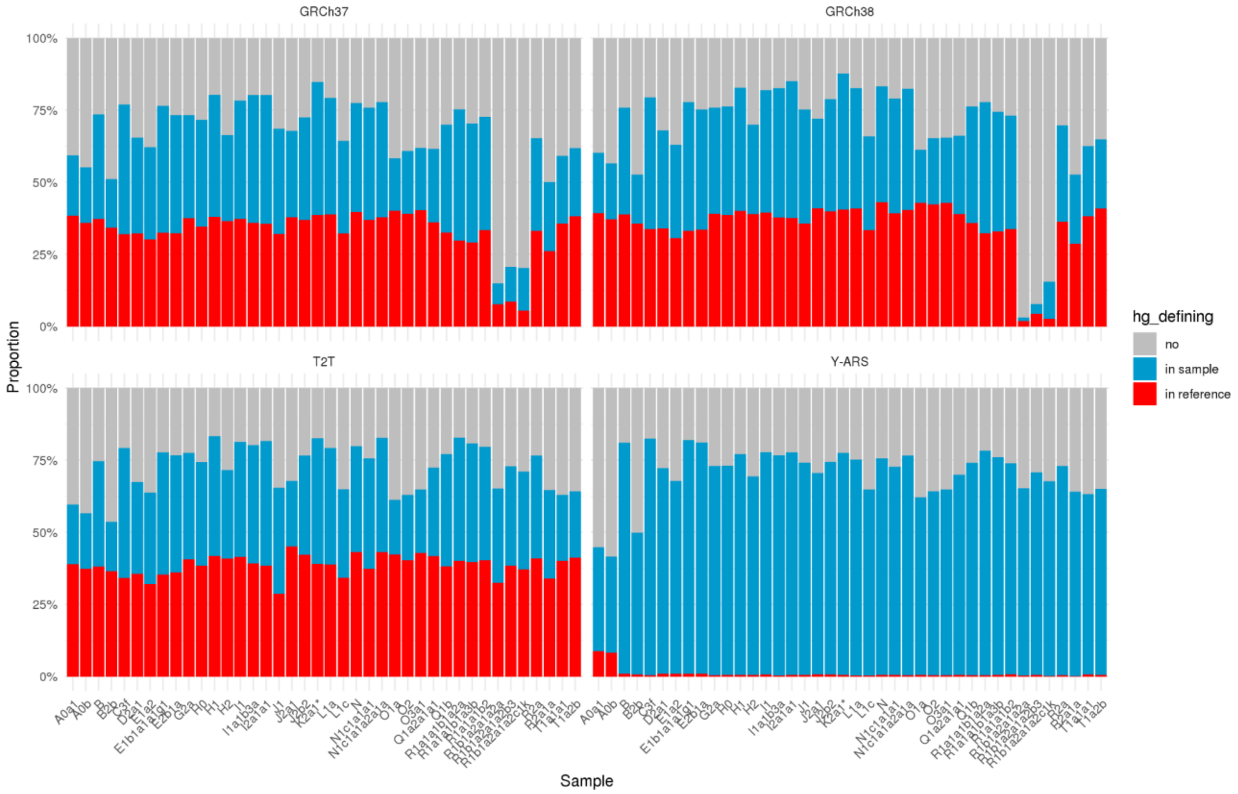


**Fig. S7** ISOGG/YFull haplogroup-defining sites in the sample (blue) and in the reference (red) across the 40 samples and each of the references. Gray bars indicate SNPs without known haplogroup annotations. Out of the haplogroup-defining sites on GRCh37, GRCh38, and T2T, approximately 53% show the haplogroup-defining allele in the sample sequence, while 47% show the haplogroup-defining allele in the reference sequence.

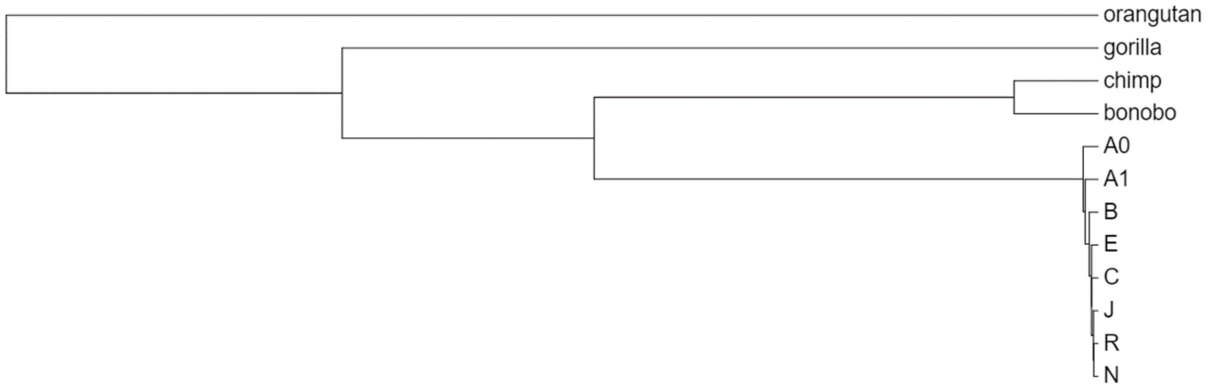


**Fig. S8** Phylogenetic tree of the four primate and eight human sequences used in the current study.

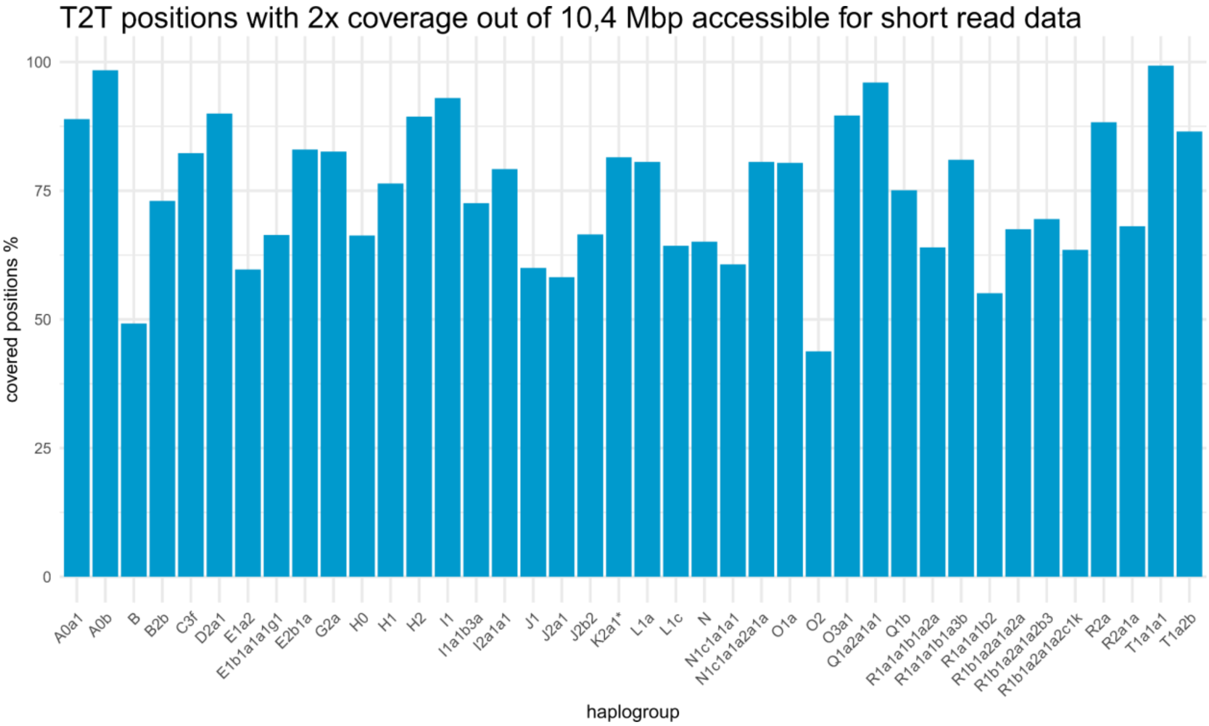


**Fig. S9** Covered positions per sample on T2T reference with coverage of at least 2x. The coverage is assessed on regions defined by Poznik et al. (2013) accessible for short-read data mapping.
